# 8-Epixanthatin Suppresses RANKL-Induced Osteoclast Differentiation via Inhibition of NF-κB and MAPK Signaling

**DOI:** 10.64898/2026.01.13.699384

**Authors:** Lifang Zhang, Vishwa Deepak

**Affiliations:** Department of Biology, College of Science, Mathematics and Technology, Wenzhou-Kean University, 88 Daxue Road, Wenzhou, Zhejiang Province 325060, China; International Frontier Interdisciplinary Research Institute (IFIRI), Wenzhou-Kean University, Wenzhou, Zhejiang Province, 325060, China; Wenzhou Municipal Key Laboratory for Applied Biomedical and Biopharmaceutical Informatics, Wenzhou-Kean University, Ouhai, Wenzhou, Zhejiang Province, 325060, China; Zhejiang Bioinformatics International Science and Technology Cooperation Center, Wenzhou-Kean University, Ouhai, Wenzhou, Zhejiang Province, 325060, China; Dorothy and George Hennings College of Science, Mathematics and Technology, Kean University, 1000 Morris Ave, Union, NJ, 07083, USA; Zhejiang-Malaysia Joint Laboratory For Rare Medicinal Resources, Wenzhou-Kean University, 88 Daxue Road, Ouhai, Wenzhou, Zhejiang Province, China

## Abstract

Osteoclast hyperactivity represents a central mechanism in pathological bone destruction, underscoring the importance of discovering novel anti-resorptive compounds. In this study, we present early-stage evidence that 8-Epixanthatin can inhibit osteoclast differentiation induced by RANKL. 8-Epixanthatin exhibited no significant cytotoxicity at the concentrations used for osteoclast differentiation studies. The compound showed concentration-dependent reductions in TRAP-positive multinucleated osteoclasts, with an IC_50_ value of 2.3 μM. Our mechanistic investigations revealed that 8-Epixanthatin interferes with RANKL-activated signaling networks, particularly NF-κB and MAPK cascades. Collectively, these observations identify 8-Epixanthatin as a promising lead structure for anti-osteoclast drug discovery.

## Introduction

Bone tissue maintains equilibrium through balanced osteoblast-mediated formation and osteoclast-driven resorption (Bolamperti et al., 2022). Pathological conditions including osteoporosis, inflammatory osteolysis, and tumor-associated bone destruction arise when this balance tilts toward excessive resorption (Terkawi et al., 2022). RANKL serves as the principal cytokine driving osteoclast differentiation, activating intracellular pathways, notably NF-κB and MAPK that induce expression of osteoclast-defining genes like NFATc1, c-Fos, TRAP, and cathepsin K (Sobacchi et al., 2025).

Available anti-resorptive therapies, primarily bisphosphonates and denosumab, effectively reduce bone loss but present tolerability concerns (Eastell et al., 2016). Extended treatment correlates with serious complications, particularly osteonecrosis of the jaw and atypical femoral fractures. These limitations fuel continued interest in discovering safer therapeutic alternatives. Natural products represent a historically productive source of drug candidates. Among these, 8-Epixanthatin (8-E), a sesquiterpene lactone derived from Xanthium species demonstrates anti-angiogenesis and antiviral properties (Romero et al., 2015). Structurally, sesquiterpene lactones contain α,β-unsaturated carbonyl groups that undergo Michael addition with nucleophilic cysteines in target proteins (Zimmermann et al., 2014). Given that both NF-κB and MAPK pathways depend on redox-sensitive cysteine residues and are essential for osteoclast differentiation, we hypothesized that 8-E might suppress osteoclastogenesis through these mechanisms.

Despite its interesting pharmacology, 8-E’s effects on osteoclast biology remain uncharacterized. This study presents initial evidence that 8-E inhibits RANKL-induced osteoclast differentiation by disrupting critical downstream signaling pathways.

## Materials and Methods

### Cell culture and osteoclast differentiation

RAW264.7 murine macrophage cells were cultured in DMEM containing 10% fetal bovine serum (FBS), 100 U/mL penicillin, and 100 μg/mL streptomycin at 37°C in a humidified atmosphere with 5% CO_2_. For osteoclast differentiation, RAW264.7 cells were differentiated into osteoclasts as described previously (Deepak et al., 2022).

Briefly, cells were plated at a density of 3 × 10^3^ cells per well in 96-well plates and stimulated with recombinant murine RANKL (30 ng/mL, R&D Systems) for 5 days, with medium changes every 2 days. 8-Epixanthatin (#HY-137974, MedChemExpress) was added at the initiation of RANKL stimulation and maintained throughout the culture period. Final DMSO concentration did not exceed 0.1% in all experiments.

### Cell viability assay

RAW264.7 cells were seeded at 1 × 10^4^ cells per well in 96-well plates and treated with indicated concentrations of 8-E (0.1-20 μM) for 48 hours. Cell viability was assessed using the CCK-8 assay (#G1613, Servicebio, Hubei, China) as per manufacturer’s instructions. Cell viability was expressed as percentage relative to vehicle-treated control cells.

### TRAP staining and quantification

After 5 days of culture, cells were fixed with 4% paraformaldehyde for 15 minutes at room temperature and stained for TRAP activity using a leukocyte acid phosphatase staining kit (#G1050, Servicebio, Hubei, China) according to manufacturer’s protocol. TRAP-positive multinucleated cells containing three or more nuclei were considered mature osteoclasts and were counted under a microscope (Keyence Microscope #BZ-X810) at 20× magnification as described previously (Rantlha et al., 2017). For each experimental condition, at least five wells were analyzed, and total osteoclasts per well were quantified. Representative images were captured at 20× magnification using a digital camera mounted on the microscope.

### Western blot analysis

RAW264.7 cells were seeded in 10-cm dishes and cultured overnight. Cells were serum-starved for 4 hours in DMEM containing 1% FBS, then pretreated with 8-E (10 μM) or vehicle (DMSO) for 2 hours. Following pretreatment, cells were stimulated with RANKL (50 ng/mL) for 15 minutes. Cells were lysed in RIPA buffer containing protease and phosphatase inhibitors. Protein concentrations were determined using the BCA assay. Equal amounts of protein (30 μg) were resolved by 10% SDS-PAGE and transferred to PVDF membranes as described elsewhere (Deepak et al., 2016). Membranes were blocked with 3% BSA in TBST for 1 hour at room temperature, then incubated overnight at 4°C with primary antibodies (Servicebio, Hubei, China) against phospho-p65 (Ser536) (#GB113882), p65(#GB11997), phospho-ERK1/2 (Thr202/Tyr204) (GB113492), ERK1/2 (#ZB12087), phospho-JNK (Thr183/Tyr185) (#GB12018), JNK (#GB114321), phospho-p38 (Thr180/Tyr182) (#GB153380), p38 (#GB154685) (1:1000 dilution), and GAPDH (#ZB15004-HRP, Servicebio, Hubei, China, 1:3000 dilution). After washing, membranes were incubated with HRP-conjugated secondary antibodies (1:5000) for 1 hour at room temperature. Protein bands were visualized using enhanced chemiluminescence (ECL) reagent and imaged using a ChemiDoc imaging system (BioRad). Densitometric analysis was performed using ImageJ software (NIH). Phosphorylated protein levels were normalized to corresponding total protein levels.

### Quantitative real-time PCR analysis

Total RNA was extracted from RAW264.7 cells after 5 days of RANKL stimulation (30 ng/mL) with or without 8-E (2.5 μM) using total-RNA isolation Kit (#G3640, Servicebio, Hubei, China) according to manufacturer’s protocol. RNA concentration and purity were determined by spectrophotometry. First-strand cDNA was synthesized from 1 μg total RNA using a reverse transcription kit (#G3333, Servicebio, Hubei, China). Quantitative real-time PCR was performed using SYBR Green Master Mix (#G3320, Servicebio, Hubei, China) on LightCycler® 96 (Roche). The primer sequences used are available on request. Relative gene expression was calculated using the 2^(-ΔΔCt) method with Gapdh as the internal control.

### Statistical analysis

Data are expressed as mean ± standard deviation (SD) of at least three independent experiments. Statistical significance was determined by unpaired Student’s t-test using GraphPad Prism 9.0 software. A *P*-value of <0.05 was considered statistically significant. \**P*<0.05, \*\**P*<0.01, \*\*\**P*<0.001, \*\*\*\**P* <0.0001 compared to RANKL-treated group.

## Results

### 8-Epixanthatin exhibits minimal cytotoxicity in osteoclast precursor cells

The chemical structure of 8-E, featuring the characteristic α,β-unsaturated carbonyl moiety of sesquiterpene lactones, is shown in Figure 1A. RAW264.7 cells were treated with increasing concentrations of 8-E (0.1-20 μM) for 48 hours. Cell viability remained ∼99% at concentrations up to 10 μM, with a modest reduction to approximately 81% observed only at 20 μM (Figure 1B). These results indicated that concentrations up to 10 μM could be used for subsequent experiments without confounding effects of cytotoxicity.

**Figure 1.**
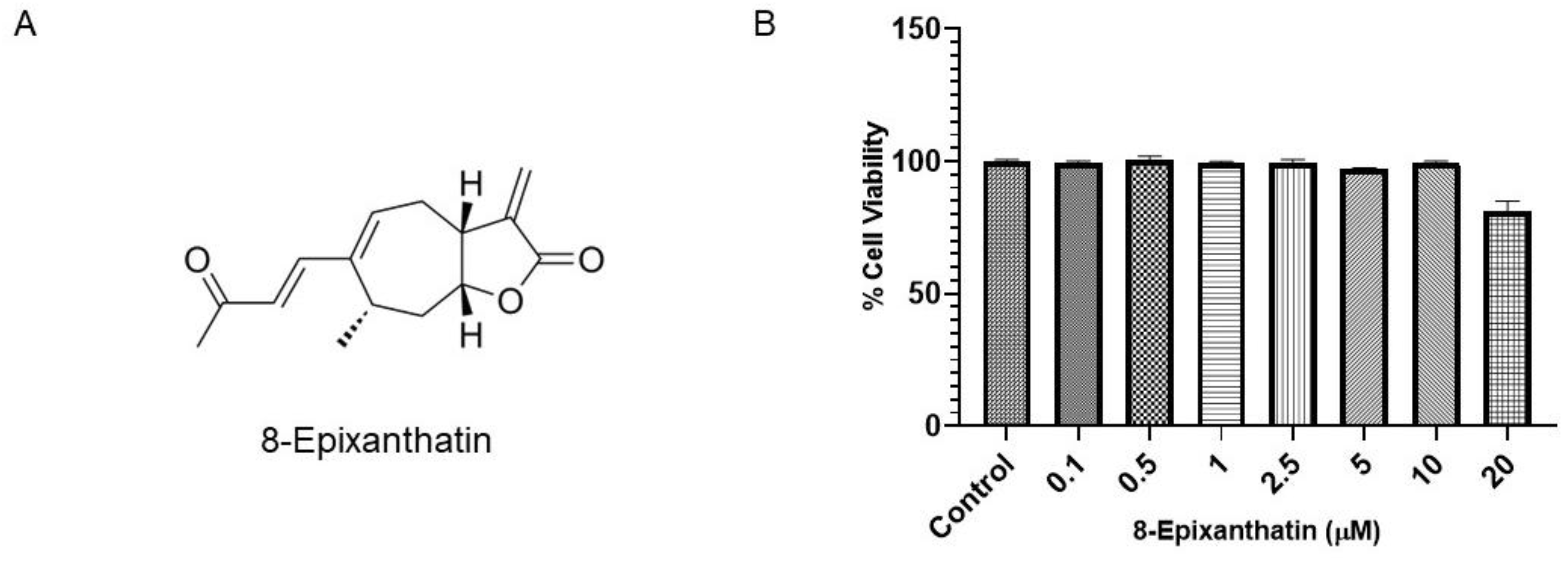
Structure and cytotoxicity profile of 8-Epixanthatin. (A) Chemical structure of 8-Epixanthatin showing the characteristic α,β-unsaturated carbonyl moiety of sesquiterpene lactones. (B) RAW264.7 cells were treated with indicated concentrations of 8-Epixanthatin (0.1-20 μM) for 48 hours, and cell viability was assessed by CCK-8 assay. Data are presented as mean ± SD from three independent experiments performed in triplicate. Cell viability remained >99% at concentrations ≤10 μM.

### 8-Epixanthatin potently inhibits RANKL-induced osteoclastogenesis

To evaluate the effect of 8-E on osteoclast differentiation, RAW264.7 cells were stimulated with RANKL in the presence of varying concentrations of the compound (0.1-10 μM). Dose-response analysis revealed concentration-dependent suppression of TRAP-activity (Figure 2A). The compound exhibited potent anti-osteoclastogenic activity with an IC_50_ of 2.3 μM, indicating that 8-E inhibits osteoclast formation in the low micromolar range. Representative TRAP staining images demonstrated abundant large, multinucleated TRAP-positive osteoclasts (appearing as dark purple cells) in RANKL-stimulated cultures, whereas treatment with 2.5 μM 8-E markedly reduced both osteoclast number and size (Figure 2B). Vehicle (DMEM) control cultures showed only scattered mononuclear cells with very few osteoclasts formed. Quantitative analysis confirmed a significant reduction in the number of TRAP-positive multinucleated cells (≥3 nuclei) from 168 ± 19 osteoclasts per well in RANKL-treated cultures to 53 ± 6 osteoclasts per well following treatment with 2.5 μM 8-E (*P*<0.0001, representing approximately 69% inhibition) (Figure 2C). These findings demonstrate that 8-E potently inhibits RANKL-induced osteoclast differentiation at concentrations well below those causing cytotoxicity.

**Figure 2.**
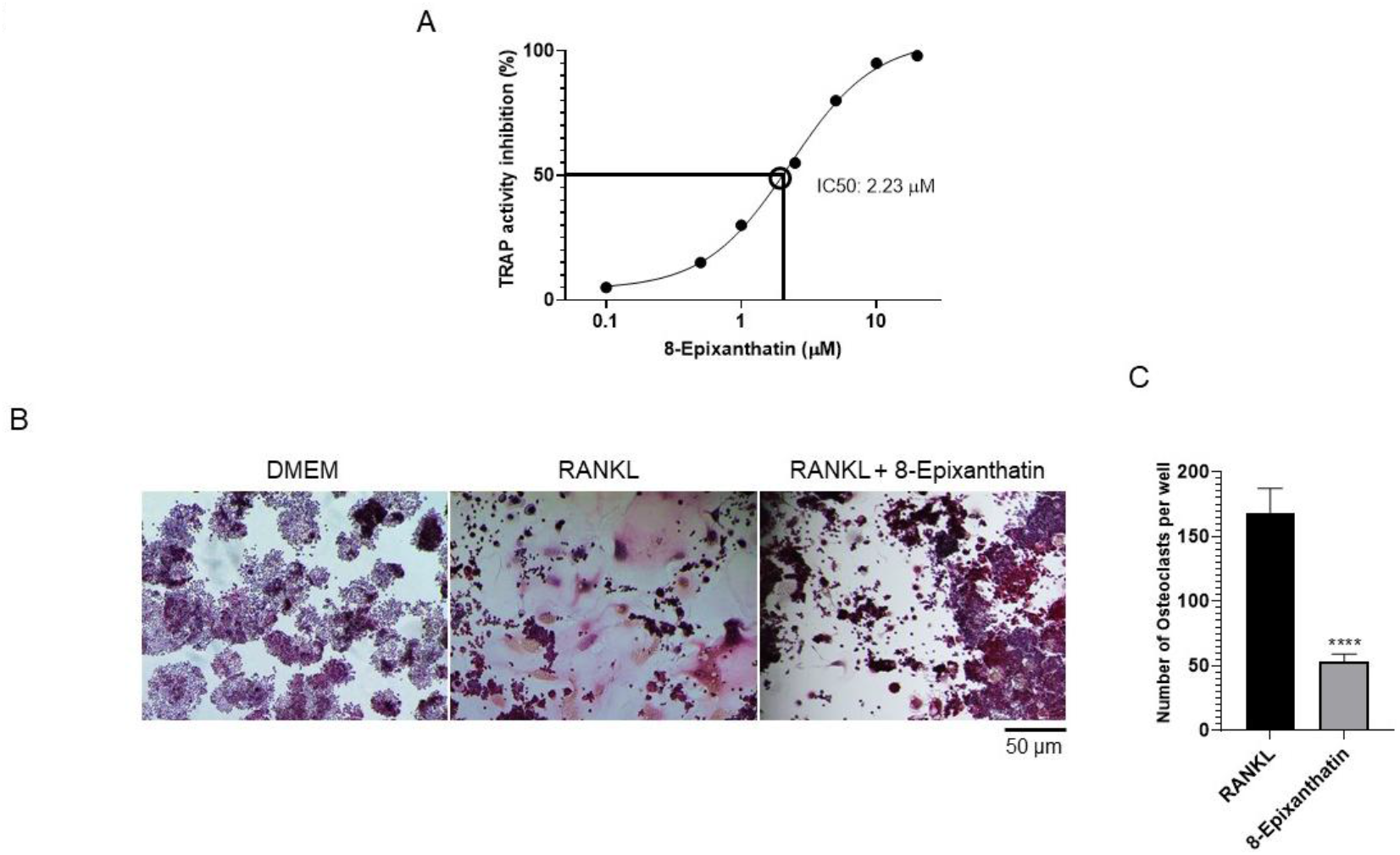
8-Epixanthatin potently inhibits RANKL-induced osteoclastogenesis in a dose-dependent manner. (A) Dose-response curve showing inhibition of osteoclast formation by 8-Epixanthatin. RAW264.7 cells were stimulated with RANKL (30 ng/mL) in the presence of increasing concentrations of 8-Epixanthatin (0.1-10 μM) for 5 days. Cells were assayed for TRAP activity (IC_50_ = 2.3 μM). Data are mean ± SD (n=3 independent experiments). (B) Representative images of TRAP-stained cultures showing osteoclast formation. Left panel: DMEM vehicle control; Middle panel: RANKL (30 ng/mL) stimulation showing numerous large, multinucleated TRAP-positive osteoclasts (dark purple cells); Right panel: RANKL + 8-Epixanthatin (2.5 μM) showing markedly reduced osteoclast number and size. Scale bar = 50 μm. Original magnification 20×. (C) Quantification of TRAP-positive multinucleated osteoclasts (≥3 nuclei) per well from cultures shown in panel B. Data are mean ± SD from three independent experiments. \*\*\*\**P*<0.0001 compared to RANKL alone by Student’s t-test.

### 8-Epixanthatin suppresses RANKL-induced NF-κB and MAPK signaling and inhibits expression of osteoclast marker genes

To elucidate the molecular mechanisms underlying 8-E’s anti-osteoclastogenic effect, we examined RANKL-activated signaling pathways. RANKL stimulation induced robust phosphorylation of NF-κB p65 at Ser536, which was reduced by approximately 30% following pretreatment with 10 μM 8-E, while total p65 levels remained unchanged (Figure 3A). Similarly, 8-E substantially suppressed RANKL-induced phosphorylation of all three MAPK family members: ERK1/2 (∼70% reduction), JNK (∼48% reduction), and p38 (∼63% reduction), without affecting total kinase expression (Figure 3A).

**Figure 3.**
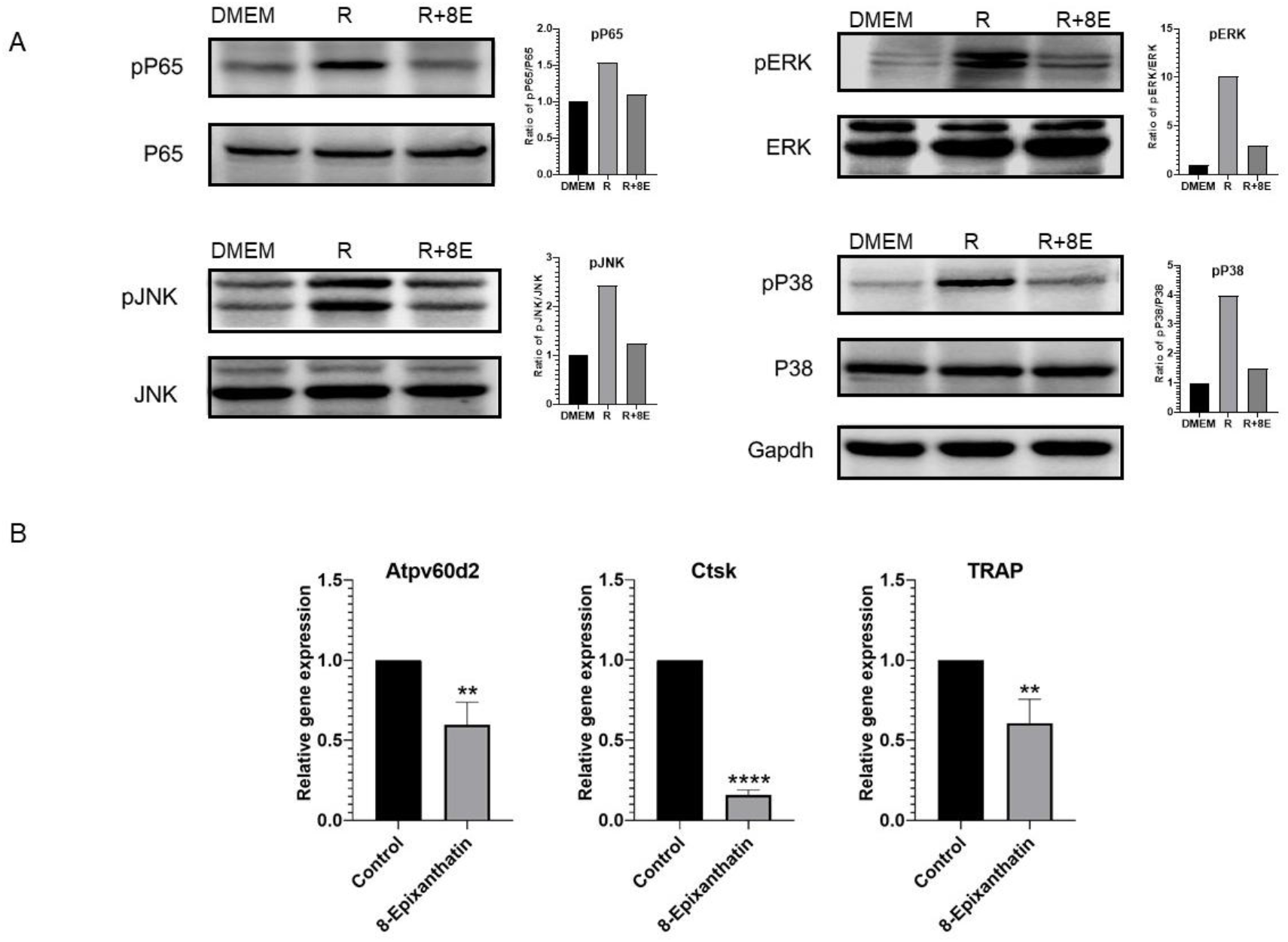
8-Epixanthatin suppresses RANKL-induced activation of NF-κB and MAPK signaling pathways and inhibits expression of osteoclast marker genes. (A) RAW264.7 cells were pretreated with 8-Epixanthatin (10 μM, 8E) or vehicle (DMSO) for 2 hours, then stimulated with RANKL (50 ng/mL) for 15 minutes. DMEM = unstimulated control. Cell lysates were analyzed by Western blot for phosphorylated and total levels of p65, ERK1/2, JNK, and p38. GAPDH served as loading control. Representative Western blots are shown with corresponding densitometric quantification in bar graphs. Phosphorylated protein levels were normalized to corresponding total protein levels. (B) RAW264.7 cells were treated with RANKL (30 ng/mL) in the presence or absence of 8-Epixanthatin (2.5 μM) for 5 days. Expression of osteoclast marker genes *Atp6v0d2, cathepsin K* (Ctsk), and *TRAP* was analyzed by quantitative RT-PCR. Data are mean ± SD from three independent experiments. **P<0.01, ****P<0.0001 compared to RANKL alone. Statistical comparisons were performed using unpaired Student’s t-test.

**Figure 4.**
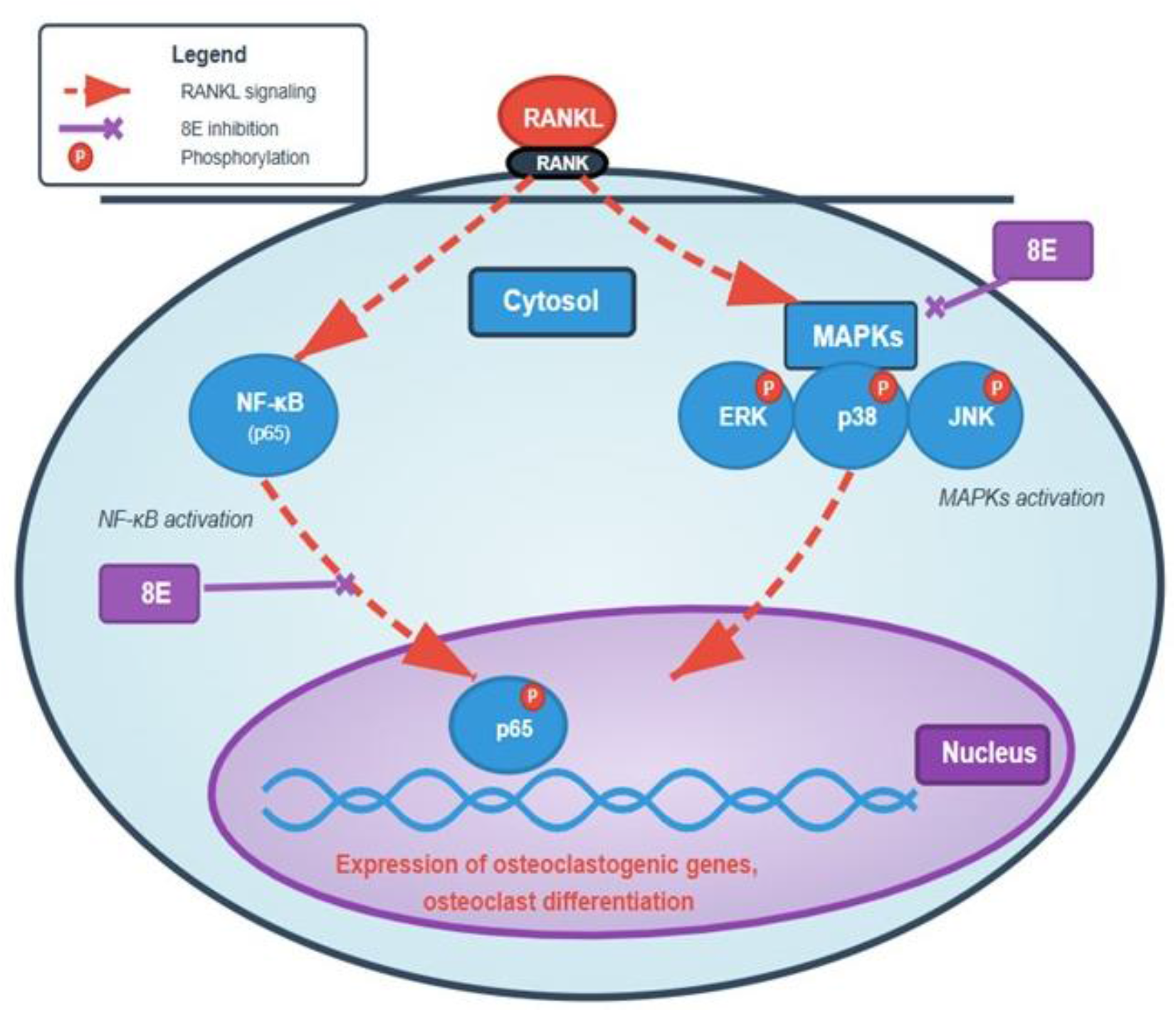
Proposed mechanism of 8-Epixanthatin inhibition of RANKL-induced osteoclastogenesis. Schematic diagram illustrating that 8-Epixanthatin (8E) suppresses RANKL-induced osteoclast differentiation by blocking NF-κB and MAPK (ERK, JNK, p38) pathway activation downstream of RANK receptor signaling. Inhibition of these pathways prevents expression of osteoclastogenic genes required for osteoclast differentiation and function.

To confirm transcriptional suppression, we examined expression of osteoclast marker genes. 8-E (2.5 μM) significantly reduced RANKL-induced expression of *Atp6v0d2* (∼40%, P<0.01), *cathepsin K* (∼85%, P<0.0001), and *TRAP* (∼40%, P<0.01) (Figure 3B). These results demonstrate that 8-E targets multiple RANKL-responsive signaling cascades and downstream gene expression essential for osteoclast differentiation.

## Discussion

This study provides preliminary evidence that 8-Epixanthatin (8-E), a natural sesquiterpene lactone derived from *Xanthium* species, potently inhibits osteoclast differentiation by targeting key RANKL-responsive signaling pathways. The compound exhibited an IC_50_ of 2.3 μM for suppression of osteoclastogenesis, which lies within a therapeutically relevant range and compares favorably with other natural product-derived osteoclast inhibitors reported in the literature. Importantly, this anti-osteoclastogenic activity was achieved without cytotoxicity at effective concentrations, supporting the potential of 8-E as a lead compound for anti-resorptive drug development.

The inhibition of both NF-κB and MAPK pathways by 8-E is mechanistically significant, as these pathways function cooperatively to drive osteoclast differentiation. NF-κB activation is required for the initial induction of NFATc1, the master regulator of osteoclastogenesis. MAPK signaling particularly through ERK and JNK regulates induction of c-Fos and AP-1 activity. These pathways cooperate with NFATc1 to drive expression of osteoclast-specific genes including TRAP, cathepsin K, and fusion-related molecules (Takayanagi, 2021). These transcriptional programs converge to promote expression of osteoclast-specific genes essential for differentiation and multinucleation. The simultaneous suppression of multiple RANKL-responsive signaling cascades by 8-E likely contributes to its potent inhibitory effect at low micromolar concentrations. Notably, the parent compound Xanthatin has been previously shown to possess anti-inflammatory properties (Liu et al., 2022; Xu et al., 2024). Our novel findings extend these observations by demonstrating that 8-E, a structural analog of xanthatin, dampens RANKL-induced osteoclast differentiation, suggesting that sesquiterpene lactones from Xanthium species may represent a class of compounds with broad activity against pathological signaling in bone and inflammatory diseases.

The α,β-unsaturated carbonyl moiety characteristic of sesquiterpene lactones is known to undergo Michael addition reactions with nucleophilic cysteine residues in target proteins (Schmidt et al., 1999). This chemical property may account for the broad inhibition of RANKL-induced phosphorylation events observed in this study, as several components of the NF-κB and MAPK pathways contain functionally important cysteine residues. However, the precise molecular targets of 8-E remain to be identified. Future studies employing chemical proteomics or structure–activity relationship analyses will be required to determine whether the compound directly modifies specific signaling proteins or interferes with upstream regulatory events.

Current anti-resorptive therapies, including bisphosphonates and denosumab, are effective in reducing fracture risk but are associated with adverse effects, particularly with long-term use (Skjodt et al., 2019). In this context, small-molecule inhibitors that modulate intracellular signaling pathways may represent alternative therapeutic strategies with distinct pharmacological profiles. Several natural products have demonstrated anti-osteoclastogenic activity in vitro, and the potency of 8-E observed here suggests it compares favorably within this class.

This study has limitations inherent to its preliminary nature. Functional bone resorption assays, assessment of downstream transcriptional regulation, and in vivo efficacy studies will be necessary to fully evaluate the therapeutic potential of 8-E. In summary, our findings identify 8-Epixanthatin as a potent inhibitor of RANKL-induced osteoclastogenesis through suppression of NF-κB and MAPK signaling pathways, providing a foundation for further investigation of sesquiterpene lactones as scaffolds for anti-resorptive drug discovery.

## Author Contributions

Vishwa Deepak: Conceptualization, Study Design and Funding Acquisition. Lifang Zhang and Vishwa Deepak performed the experiments and wrote the manuscript. All authors approved the submitted version.

## Data Availability Statement

The data that support the findings of this study are available from the corresponding author upon reasonable request.

## Institutional Review Board Statement

Not applicable. This study used only the established RAW264.7 cell line and did not involve human subjects or animal procedures.

## Acknowledgments

This work was supported by WKU-ISRG grant (ISRG2023032) and International Frontier Interdisciplinary Research Institute (IFIRI-WKU) grant (KY20250604000450). We thank core facilities of Wenzhou-Kean University for the resources.

